# Temporal patterns of damage and decay kinetics of DNA retrieved from plant herbarium specimens

**DOI:** 10.1101/023135

**Authors:** Clemens L. Weiß, Verena J. Schuenemann, Jane Devos, Gautam Shirsekar, Ella Reiter, Billie A. Gould, John R. Stinchcombe, Johannes Krause, Hernán A. Burbano

**Affiliations:** Research Group for Ancient Genomics and Evolution, Department of Molecular Biology, Max Planck Institute for Developmental Biology; Institute of Archaeological Sciences, University of Tübingen; Department of Molecular Biology, Max Planck Institute for Developmental Biology; Department of Ecology and Evolutionary Biology, University of Toronto; Centre for the Analysis of Genome Evolution and Function, University of Toronto; Departments of Paleoanthropology and Archaeogenetics, Senckenberg Center for Human Evolution and Paleoenvironment, University of Tübingen; Max Planck Institute for the Science of Human History

**Author notes:** Corresponding author: Hernán A. Burbano Group Leader – Ancient Genomics and Evolution, Department of Molecular Biology, Max Planck Institute for Developmental Biology, Spemannstr. 37-39. Tuebingen, D-72076, Germany.

## Abstract

Herbaria archive a record of changes of worldwide plant biodiversity harboring millions of specimens that contain DNA suitable for genome sequencing. To profit from this resource, it is fundamental to understand in detail the process of DNA degradation in herbarium specimens. We investigated patterns of DNA fragmentation -length and base composition at breaking points-, and nucleotide misincorporation by analyzing 86 herbarium samples spanning the last 300 years using Illumina shot-gun sequencing. We found an exponential decay relationship between DNA fragmentation and time, and estimated a per nucleotide fragmentation rate of 1.66 × 10^−4^ per year, which is ten times faster than the rate estimated for fossilized bones. Additionally, we found that strand breaks occur specially before purines, and that depurination-driven DNA breakage occurs constantly through time and can to a great extent explain decreasing fragment length over time. Similar of what has been found analyzing ancient DNA from bones, we found a strong correlation between the deamination-driven accumulation of cytosine (C) to thymine (T) substitutions and time, which reinforces the importance of substitution patterns to authenticate the ancient/historical nature of DNA fragments. Accurate estimations of DNA degradation through time will allow informed decisions about laboratory and computational procedures to take advantage of the vast collection of worldwide herbarium specimens.

## Introduction

Under favorable conditions DNA fragments can survive in plant (Palmer et al. 2012) and animal tissues (Allentoft et al. 2012) for thousands of years providing a molecular record of the past. Therefore, the examination of historical genomes permits the inclusion of temporal data into evolutionary studies, which allows a more accurate inference of rates and timing of key evolutionary events, e.g. hybridization, speciation, or mutation. During the last decade, and thanks to the advent of high-throughput sequencing (HTS), the study of DNA retrieved from historic samples has changed our views on different fields ranging from human evolution (Green et al. 2010; Rasmussen et al. 2010; Reich et al. 2010; Meyer et al. 2012) to the emergence and re-emergence of both plant (Martin et al. 2013; Yoshida et al. 2013; Smith et al. 2014) and human pathogens (Bos et al. 2011; Schuenemann et al. 2013; Bos et al. 2014). The vast majority of ancient DNA (aDNA) studies have focused on animal remains, especially fossilized bones and teeth, whereas plant remains have received less attention (Shapiro and Hofreiter 2014) despite the abundance of historic plant specimens.

In general, DNA retrieved from historic specimens comes in small amounts and is a mixture of endogenous and microbial DNA that either was present *pre-mortem* or colonized the tissue *post-mortem* (Green et al. 2006; Poinar et al. 2006; Zaremba-Niedzwiedzka and Andersson 2013). Ancient DNA comes in small fragment sizes (Paabo 1989) and holds various modifications that distinguish it from DNA extracted from fresh tissue (Dabney et al. 2013b). It has been shown with *in vitro* experiments that DNA fragmentation is partially driven by spontaneous depurination and subsequent hydrolysis of the DNA backbone (Lindahl and Andersson 1972; Lindahl and Nyberg 1972). The typical sign of depurination, which is the excess of both adenine (A) and guanine (G) before DNA breaking points, has been detected by HTS in libraries constructed from aDNA (Briggs et al. 2007). DNA degradation is additionally marked by an increase of cytosine (C) to thymine (T) substitutions towards the end of aDNA fragments. This has been interpreted as a result of spontaneous deamination of C residues to uracils (U) that are read as T by the polymerase and occur in higher proportion in single-stranded DNA overhangs (Briggs et al. 2007; Brotherton et al. 2007). A biochemical definition of aDNA includes all above-mentioned characteristics but does not delineate a time boundary between ancient and modern DNA (Shapiro and Hofreiter 2014).

It is particularly interesting to understand quantitatively how these aDNA-associated patterns change through time, since they could be used to both authenticate DNA fragments retrieved from historic samples of different ages, and to calculate DNA decay rate based on their fragmentation patterns (Allentoft et al. 2012). Using fossilized animal remains it has been found that there is a strong negative correlation between the amount of putative deamination (excess of C to T substitutions) and the sample age (Sawyer et al. 2012). Hence, the excess of C to T substitutions has been repeatedly used as criteria of authenticity in aDNA studies; it has proved to be particularly useful in the study of ancient modern human remains (Krause et al. 2010; Prufer and Meyer 2015). The correlation between other aDNA-associated patterns and sample age is weaker (Sawyer et al. 2012), which could be a consequence of the very different environmental conditions in which fossils were preserved, collected, processed and stored. It was suggested that conditions such as temperature, pH, and humidity, among many others, affect DNA stability (Lindahl 1993), but deamination rate seems to be resilient to variation in these environmental conditions (Sawyer et al. 2012). To reduce the effect of environmental variation on DNA degradation, a more spatially constrained sample of extinct moas, a flightless bird endemic to New Zealand, has been studied (Allentoft et al. 2012). Allentoft et al. (2012) work calculated the long-term DNA decay rate in bone tissue, which could be used to estimate DNA half-life and consequently to put a boundary on how far back in the past DNA could be theoretically retrieved. Because most of the bone samples in Allentoft et al. (2012) were analyzed only by quantitative PCR and not by HTS, it was not possible to investigate damage patterns of the original molecule ends, since the amplifiable template starts 3’ after the annealed primer. Therefore they could not look at how the signals left by deamination or depurination correlate with time in a spatially constrained sample.

Although herbaria are ubiquitous in natural history museums and harbor millions of plant specimens, they have not been extensively sampled for genetic analyses, particularly using library-based methods coupled with HTS. These methods are ideal to recover small fragments typical for aDNA, since adaptor DNA molecules can be efficiently ligated to fragments of short size. By contrast direct PCR tends to preferentially amplify longer DNA fragments from a distribution of aDNA molecules, since primers need to correctly anneal before amplification. Since herbaria contain time snapshots of global biodiversity and could be informative to address a broad spectrum of biological questions, it is fundamental to understand how DNA survives in this type of specimens. It is highly advantageous that herbaria samples are prepared and stored using standardized procedures, which reduces the amount of environmental variation among herbaria samples compared with fossilized bones. Consequently, herbarium samples are ideal to study the temporal patterns of damage and decay kinetics of DNA. In this study, we analyzed 86 herbarium samples collected over the last 300 years using library based-based methods coupled with HTS and produce for the first time an in depth description of aDNA associated patterns and its dynamics through time. Additionally, we use the power of multiple DNA sequencing libraries to calculate DNA decay rate and half-life in plant desiccated tissue.

## Results

### DNA fragmentation

We used a group of multiple species herbarium samples from the 18^th^, 19^th^ and 20^th^ centuries and also freshly prepared (< 1 year old) herbarium samples of *Arabidopsis thaliana* dried using a wooden press (Table 1 and Supplementary Table 1). From here on we will refer to these groups as historic and modern herbarium, respectively. A fraction of historic herbarium samples from both *Solanum tuberosum* and *Solanum lycopersicum* have lesions compatible with *Phytophthora infestans* infection and have been previously studied (Yoshida et al. 2013). All samples were paired-end sequenced using different models of the Illumina platform (Supplementary Table 2). Since we expect that DNA retrieved from historic samples will be highly fragmented and therefore shorter than the sequencing read length (101-150 bp), for most of the fragments a part of the library adapter will be sequenced. Additionally we expect that also a fragment of the molecule will be covered by both the forward and reverse read. After adapter trimming forward and reverse reads were merged requiring an overlap of 10 base pairs (bp) between them. We were able to merge the vast majority of the reads (83%-99%) from historic herbarium samples, whereas only a very small fraction of reads (18%-40%) could be merged from modern herbarium samples, due to the presence of much longer DNA fragments (Supplementary Table 2). In the historic herbarium samples, merged reads reflect the original length of the molecules to which adapters were ligated during library preparation. For the modern herbarium samples, we estimated the original length of the insert after paired-end mapping. For this group of samples, the mean of the fragment length distribution corresponded to the fragment size intended during sonication (400 bp) and the merged reads were located at the left tail of the fragment length distribution (Supplementary Fig. 1). For all further analysis correlating DNA properties with time we used only merged reads from historic herbarium samples.

**Table 1.** Type and number of herbarium samples.

| Type of Sample | Species | Number of samples | Collection year (range) | Number of infected samples<sup>*</sup> |
| --- | --- | --- | --- | --- |
| Historic | *Arabidopsis thaliana* | 54 | 1863-1993 | - |
| Historic | *Solanum tuberosum* | 12 | 1845-1896 | 12 |
| Historic | *Solanum lycopersicum* | 5 | 1737-1876 | 2 |
| Historic | Total | 71 | 1737-1993 | 14 |
| Modern | *Arabidopsis thaliana* | 15 | 2014 | - |

<sup>\*</sup>Samples with lesions compatible with *Phytophthora infestans* lesions

The high percentage of merged reads (83%-99%) shows that DNA retrieved from historic herbarium samples is indeed highly fragmented (median 44-87 bp) (Fig. 1A and Supplementary Table S2). The distribution of fragment lengths of merged reads is not normally distributed and could be better described by a log-Normal distribution (Figure 1A). To evaluate the correlation between fragment lengths with the collection year of each sample we chose the log-mean value of a fitted log-Normal distribution. The log-mean describes better the distribution of fragment lengths than other measures of central tendency, e.g. the mean, and allows performing a linear regression in an otherwise exponential relationship. The regression between the log-mean fragment length and the sample collection year was statistically significant (R^2^ = 0.2; P = 6.33 * 10^−5^). For visualization we plotted the median of each distribution of fragment lengths on a log-scaled y-axis (Figure 1B). To check if the signal was driven only by the oldest 18^th^ century *S. lycopersicum* samples (Figure 1B), we repeated the analysis only for the *A. thaliana* samples, since the majority of samples are from this species and their collection dates are widely collection year was still significant (R^2^ = 0.175; P = 1.6 * 10^−3^), which implies that the signal arises from the whole set of herbarium specimens and is not driven only by the oldest samples. Since DNA was extracted from some herbarium specimens using CTAB and PTB extraction protocols (Kistler 2012), we evaluate the effect of these methods on the length distribution of DNA reads and found no difference between them (P = 0.75) (Supplementary Fig. 2).

**Figure 1.**
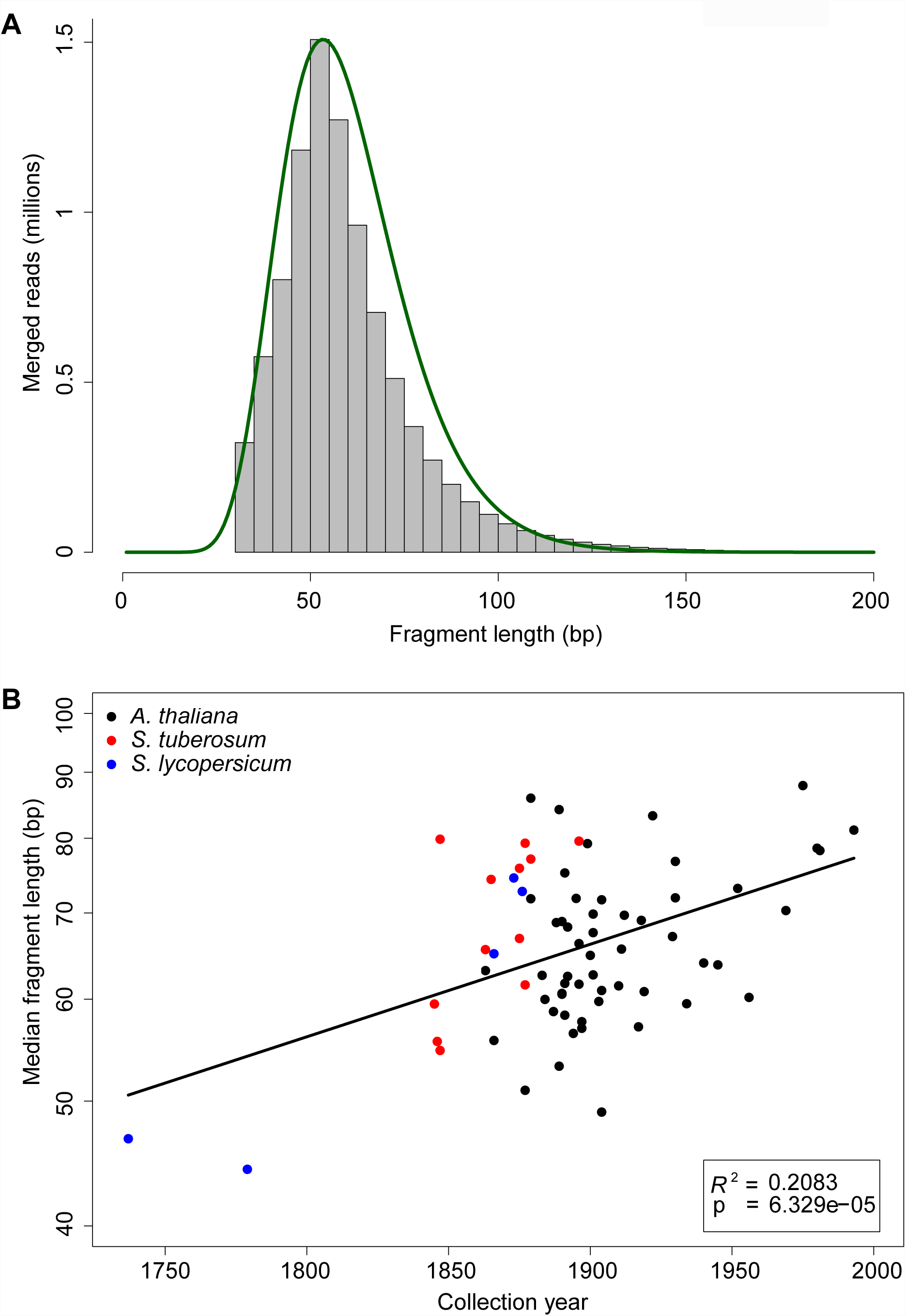
DNA fragmentation. **(A)** Distribution of fragment lengths of merged reads from *A. thaliana* sample NY1365354. The green line shows the fit between the empirical and the log normal distributions. **(B)** Median length of merged reads as a function of collection year. The line indicates the linear regression. The inset shows the regression statistics between the natural logarithm of median length and collection year. The y-axis is log scaled and shows therefore that the correlation is exponential.

### DNA break points

It has been shown based on in vitro assays with modern DNA that fragmentation is driven by depurination followed by hydrolysis of the DNA backbone (Lindahl and Andersson 1972; Lindahl and Nyberg 1972). Using reads mapped to their respective reference genome, it is possible to analyze the genomic nucleotide context surrounding the ends of the DNA fragments, and thus look indirectly at DNA break points. It has been found that there is an excess of purines (both adenine and guanine) at aDNA break points, which has been interpreted as a sign of depurination (Briggs et al. 2007). We found an excess of purine frequency (both adenine and guanine) in DNA retrieved from herbarium samples at position -1 (5’ end) (Fig. 2A). We calculated the relative enrichment in purine frequency of both adenine and guanine at position -1 compared with position -5. We then correlate these signatures of depurination with the collection year of the sample. Neither adenine (Fig. 2B) nor guanine (Fig. 2C) relative enrichment showed a significant correlation with collection year. Additionally we did not find a difference between the average relative enrichment of adenine when compared with guanine (Fig. 2B-C). Since chloroplast genomes occurred in high copy number and its circular structure differ enormously from the nuclear genome, we performed the same analysis independently for chloroplast-derived reads. We again found purine enrichment at position -1, and no correlation between the relative enrichment of purines and collection year. There were no significant differences between nuclear- and chloroplast-derived reads (P(adenine) = 0.34; P(guanine) = 0.7) (Supplementary Figure 3).

**Figure 2.**
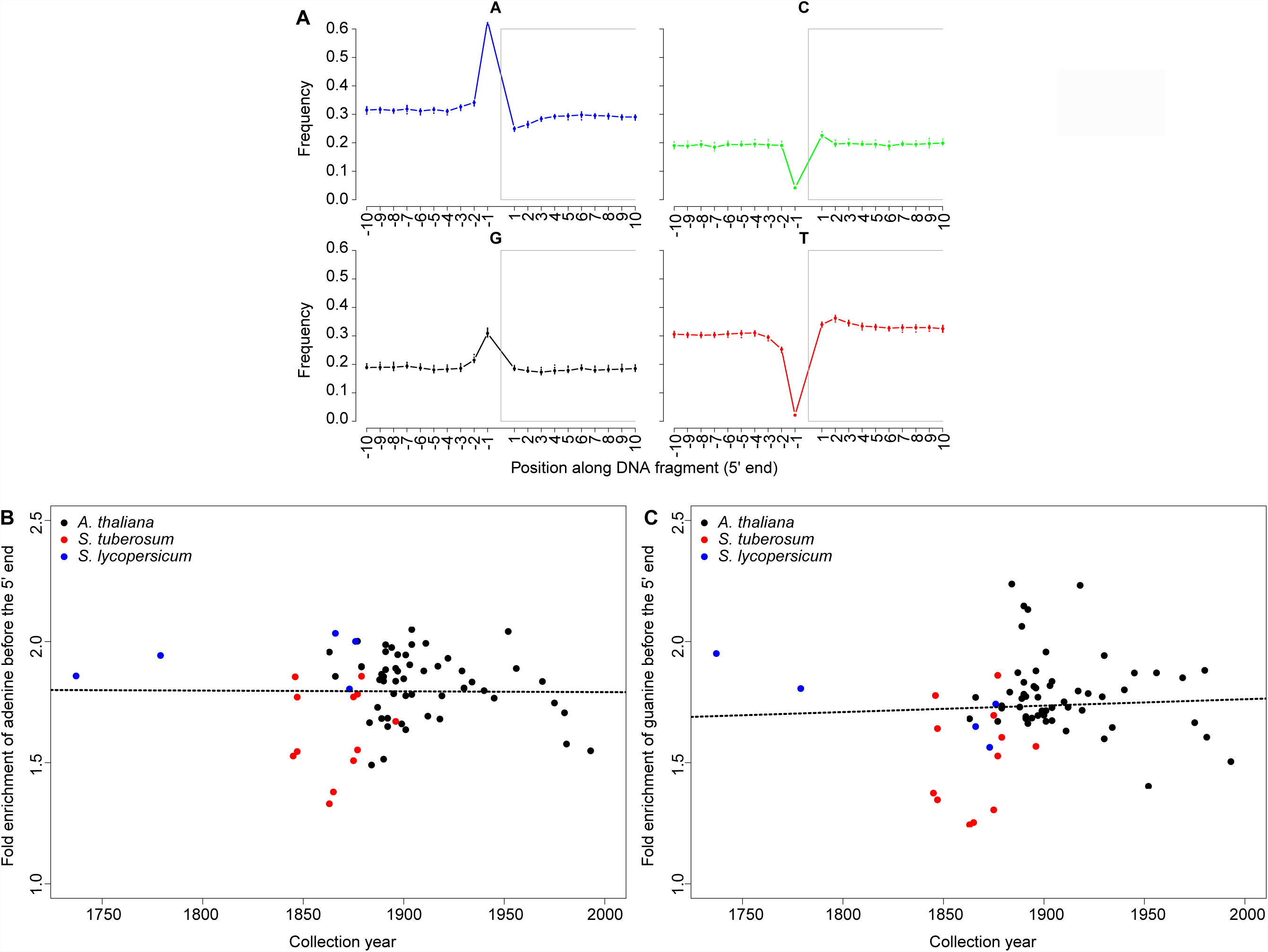
DNA breaking points. **(A)** Base composition of the first 10 nucleotides of the reads and of 10 nucleotides upstream genomic context in *A. thaliana* sample NY1365354 (after read mapping). The grey boxes separate the reads (position 1 to 10) from their upstream genomic context (positions -10 to -1). **(B)** Relative enrichment of adenine at 5’ end (position -1 compared with position -5) as a function of collection year **(C)** Relative enrichment of guanine at 5’ end (position -1 compared with position -5) as a function of collection year. In both B and C the dotted lines show the linear regression.

### DNA decay rate

For each library we used the fragment lengths of merged reads to calculate the rate of DNA decay (Allentoft et al. 2012), i.e. the pace at which bonds are broken in the DNA back bone in a per year basis. The length distribution in aDNA libraries shows an exponential decline as the result of random fragmentation (Fig. 3A) (Deagle et al. 2006). After logarithmic transformation of the fragment length frequencies, the exponential decline can be described by a linear function with slope *λ* (lambda), which corresponds to the damage fraction per site (Fig. 3B). Damage should be interpreted here as DNA bond breaking. To get the overall decay rate for all herbarium samples, we analyzed the relationship between lambda and the age of each sample and found a linear relation. The slope corresponds to the overall per nucleotide decay rate *k* = 1.66 * 10^−4^ per year (R^2^ = 0.26; P = 6.46 * 10^−6^) (Fig. 3C), which turned out to be 10 times faster than the rate estimated based on fossilized bones *k = 2.71 * 10^−5^* per site per year (Allentoft et al. 2012) (Figure 4A). We calculate the average half-life of a 100 bp fragment to be 40 years (Figure 4B).

**Figure 3.**
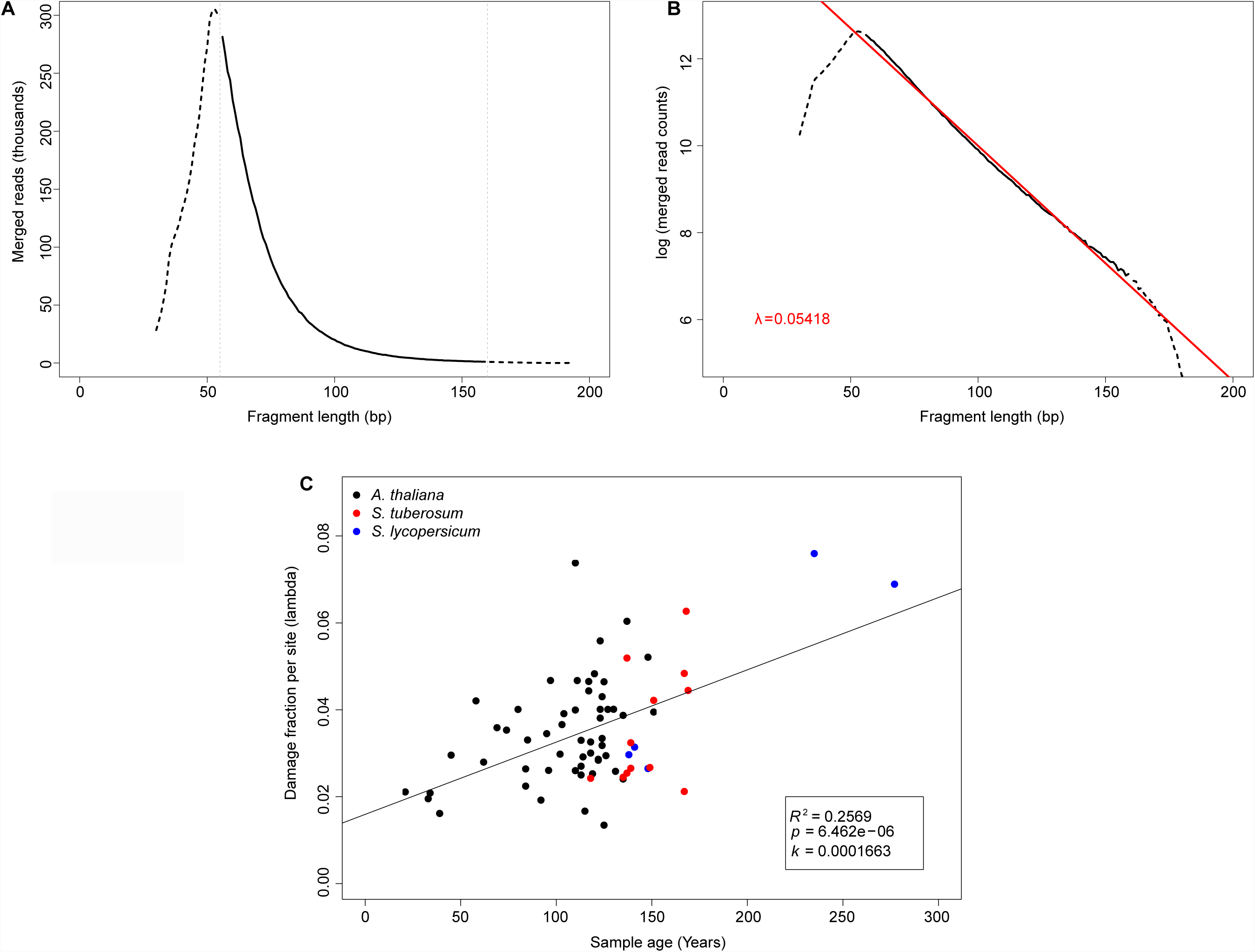
DNA fragmentation and decay rate. **(A)** Distribution of fragment lengths of merged reads from *A. thaliana* sample NY1365354. The solid line, which is surrounded by horizontal dotted lines, shows the part of the distribution that follows an exponential decline. **(B)** Distribution of fragments length for the same library using a y-axis with a logarithmic scale. The slope of the exponential part of the distribution (red line) corresponds to the damage fraction per site (lambda). **(C)** Damage fraction per site (lambda) as a function of sample age. The slope of the regression corresponds to the DNA decay rate (k) following the formula: *λ* = k * age.

**Figure 4.**
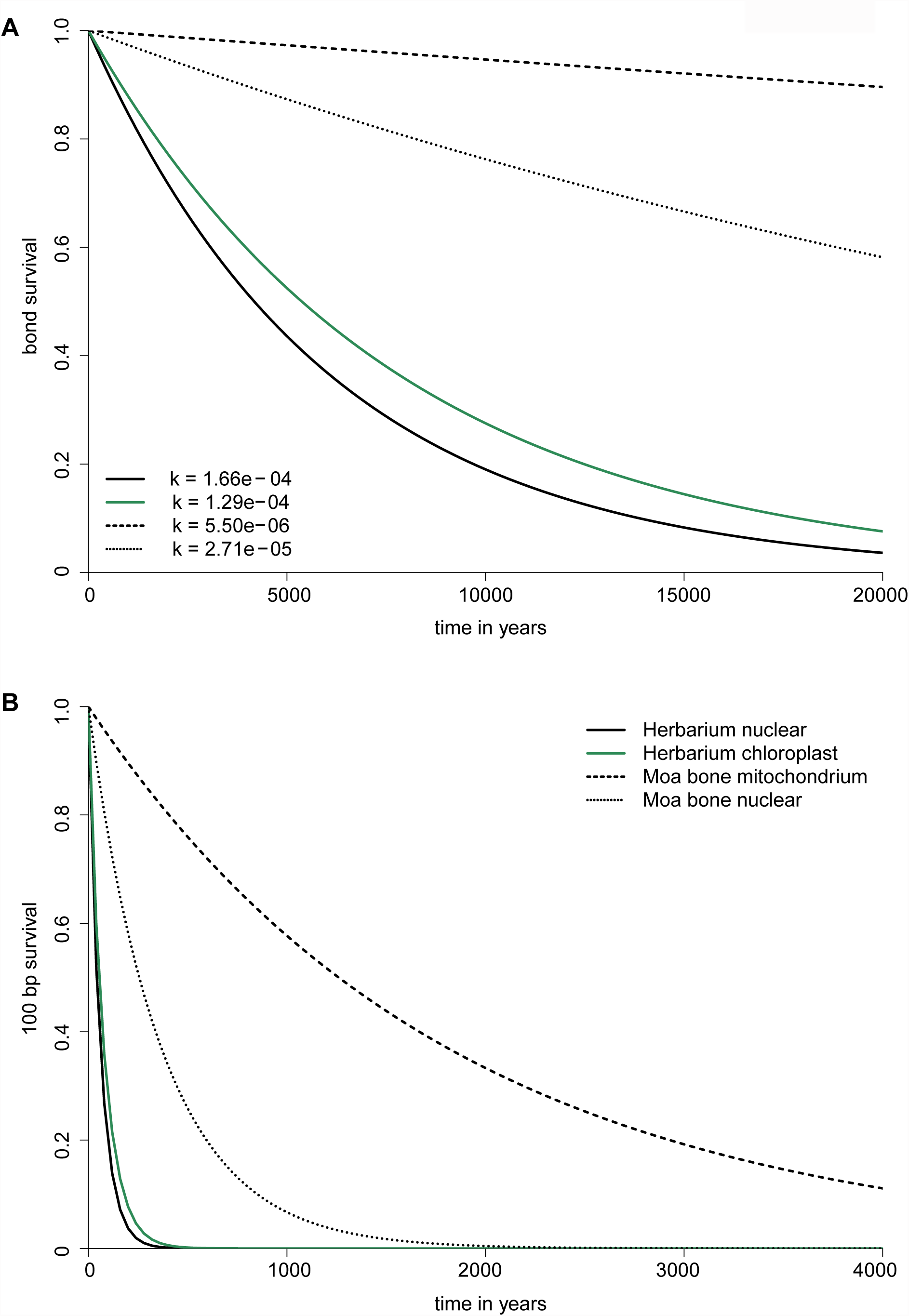
Rates of DNA decay and inferred half-life of DNA. **(A)** The survival of DNA in bone and plant desiccated tissue quantified as the survival of bond in the DNA backbone. **(B)** Estimated half-life of a 100 bp fragment using the decay rates from A.

### Nucleotide misincorporation

The most abundant miscoding lesions in aDNA are C to T substitutions, which are believed to be caused by deamination of C to U. The U is then read as T by the polymerase during sequencing (Briggs et al. 2007; Brotherton et al. 2007). The excess of C to T substitutions occurs primarily at the ends of the reads and declines exponentially inwards. We found this pattern present in all historic herbarium samples analyzed (Fig. 5A). Since the excess of C to T substitutions is more manifest at first base, we chose, as previously described (Sawyer et al. 2012), the percentage of C to T substitutions at first base as a proxy for miscoding lesions and correlate this value with the samples’ collection year (Fig. 5B). We found a very strong linear relationship between these two values, which can be shown by a linear regression among the percentage of C to T substitutions at first base and the sample collection year (R^2^ = 0.45; P = 1.44 * 10^−10^).

**Figure 5.**
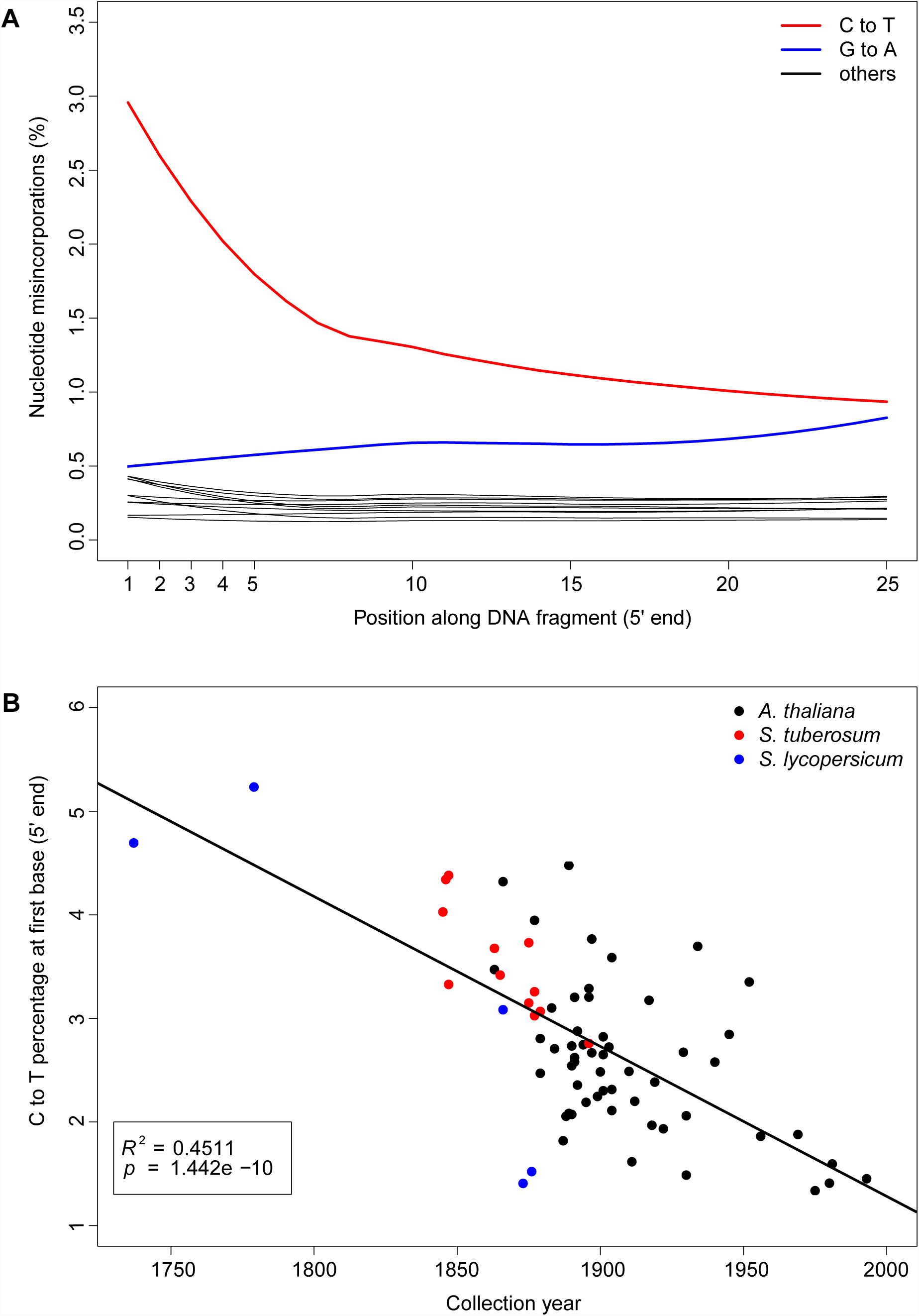
Nucleotide misincorporation. **(A)** Nucleotide misincorporation profile at 5’end of the reads of *A. thaliana* sample NY1365354. The red line shows an excess of C to T substitutions at the beginning of the read that declines exponentially inwards. **(B)** C to T percentage at first base (5′ end) as a function of collection year. The C to T percentage and the collection year have a linear relationship.

For the infected samples we also calculated the percentage of C to T substitutions in *Phytophthora infestans* derived reads at first base and found the same signature, although it was weaker than the signal found in their host plant (Supplementary Fig. 4).

### Differences between nuclear- and chloroplast-derived reads

We found that chloroplast-derived reads showed a slightly lower decay rate than the nuclear-derived reads (*k*_*chloroplast*_ = 1.29 * 10^−4^) (Fig. 6A). However the regression is weaker due to a wider spread of data points caused by the fact that less reads align to the chloroplast genome compared with the nuclear genome (R^2^ = 0.14; P = 1.2 * 10^−3^). To test if the two decay rates were different we performed an analysis of variance that showed significant effects of both sample age and origin of DNA (nuclear- or chloroplast-derived) on the rate of bond breaking (lambda) (Pr(sample age) = 4.84 * 10^−8^, Pr(DNA origin) = 0.012). However, the effect of DNA origin was very low and there was no significant interaction between sample age and DNA origin (Pr(Sample age:DNA origin) = 0.46). This indicates that the slopes of the two regressions, which correspond to the decay rate k, do not differ significantly.

**Figure 6.**
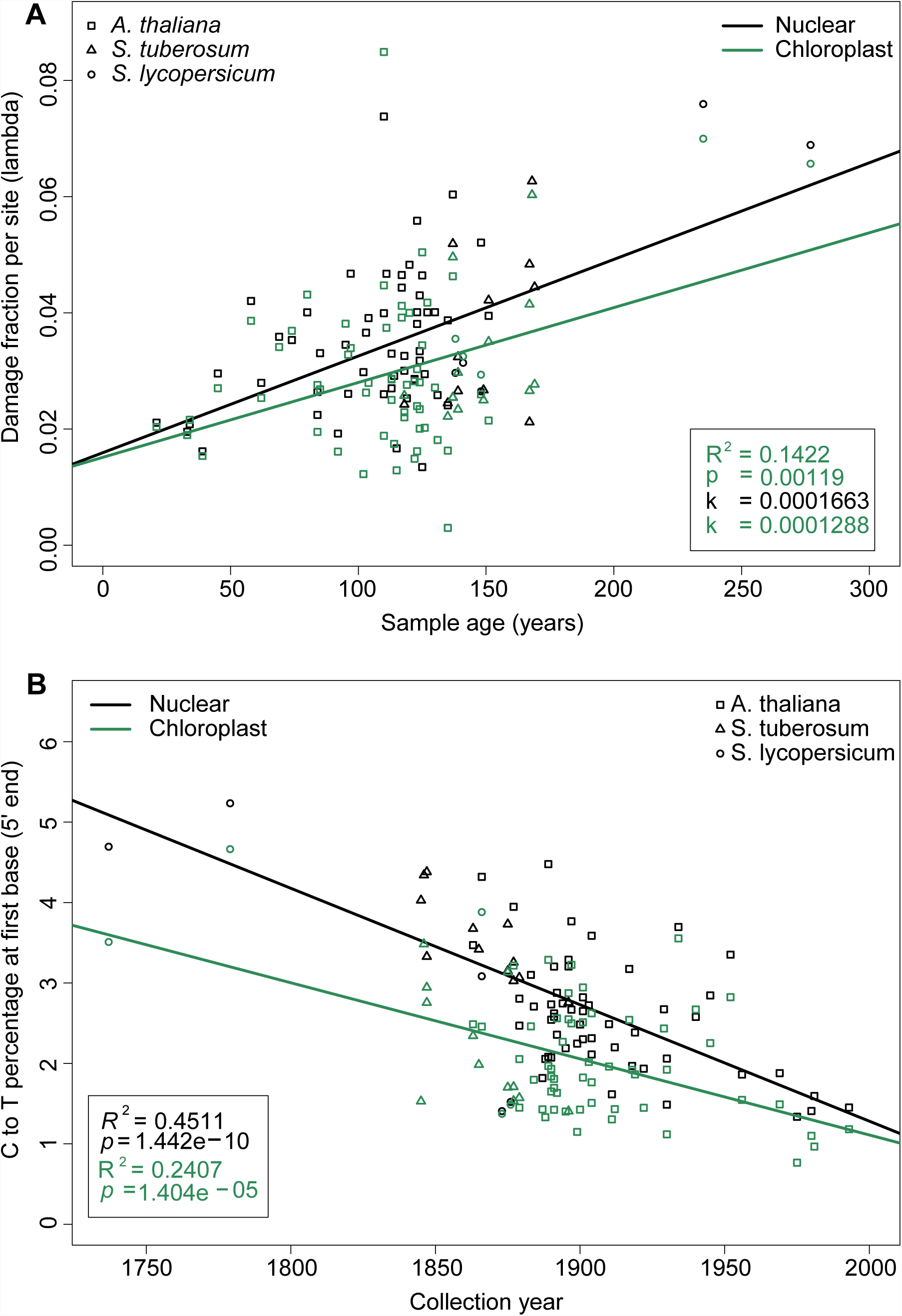
DNA decay and misincorporation in nuclear and chloroplast reads. **(A)** Damage fraction per site (lambda) as a function of sample age. **(B)** C to T frequency at first base (5 end) as a function of collection year. In both (A) and (B) data points and values for nuclear and chloroplast reads are shown in black and green font, respectively.

The chloroplast-derived reads show a lower excess of C to T substitutions than the nuclear-derived reads (Fig. 6B). As it was found in the decay rate, the data points were more spread than the nuclear-derived reads, which causes a weaker regression (R^2^ = 0.24; P = 1.4 * 10^−5^). The analysis of variance showed in this case highly significant effects of sample age and DNA origin on the percentage of deamination at first base (Pr(sample age) = 1.78 * 10^−14^, Pr(DNA origin) = 5.69 * 10^−9^). However, as it was observed in the DNA decay rate case, there was no significant interaction between sample age and DNA origin (Pr(sample age:DNA origin) = 0.075). This indicates that nuclear- and chloroplast-derived sequences differ significantly in the extent of deamination (the intersect of the regressions), but not in its rate (slope of the regressions).

## Discussion

Herbaria contain millions of dried plant specimens that provide a record of worldwide changes in biodiversity spanning five centuries. Although plants were not originally collected and stored for genetic studies, the value of these collections as source of DNA has been long recognized by plant biologists (Rogers and Bendich 1985). There are a larger number of studies that have used PCR-based approaches to survey these collections, but only a handful of endeavors have used library-based methods coupled with HTS (Martin et al. 2013; Staats et al. 2013; Yoshida et al. 2013). Since herbaria collections are an invaluable source of genetic information, it is important to investigate in detail both the properties of DNA retrieved from them, and the rate at which DNA damage takes place through time.

There are two important characteristics of our study that have to be taken into account before we compare it with previous investigations that have utilized plant and animal remains: (i) The vast majority of studies employing herbarium samples have used PCR-based approaches, which preferentially amplify long DNA fragments and survey only a small subset of the total of DNA molecules. By contrast, we have used a library-based method coupled with HTS, which allows analysis of an extensive population of DNA molecules that better represents the molecule length distribution preserved in the DNA extract; (ii) The correlation between aDNA properties and time has been investigated using fossilized animal remains that have experienced a wide range of environmental conditions (Sawyer et al. 2012). Although one study tried to minimize environmental variation using a more spatially constrained set of samples (Allentoft et al. 2012), we argue here that herbarium samples have experienced less environmental variation, since they have been collected, prepared and stored in a very standardized way, which makes them ideal to study temporal patterns of DNA damage and decay.

### DNA fragmentation

We confirmed the highly fragmented nature of DNA retrieved from herbarium samples (Staats et al. 2013; Yoshida et al. 2013), which is more accurately described by a log-Normal distribution (Fig. 1A). The DNA fragmentation is comparable with the level found in fossilized animal remains that are several hundreds or even thousands of years old (Sawyer et al. 2012), although our samples are at most 278 years old. In contrast to fossilized animal remains that showed no correlation between fragment length and age (Sawyer et al. 2012), we found a significant exponential relation between fragment length and collection year, where more recent samples have longer DNA fragments (Fig. 1B). The lower levels of environmental variation experienced by herbarium samples relative to fossilized animal remains could have increased the signal to noise ratio allowing the detection of the relation between time and DNA fragmentation.

Since it has been shown *in vitro* that DNA fragmentation is driven by hydrolytic depurination followed by ß-elimination (Lindahl and Andersson 1972; Lindahl and Nyberg 1972), and depurination can be inferred by examining DNA breaking points in HTS data (Briggs et al. 2007; Sawyer et al. 2012), we analyzed our sequencing libraries for an excess of purines at genomic positions surrounding sequencing reads. Both A and G were found overrepresented upstream of the 5’ end break points (Fig. 2A), but no correlation was found between the relative fold enrichment of either A or G, and collection year (Fig. 2B-C), which implies that the contribution of depurination to DNA breakage does not change through time. Only a very weak negative correlation between age and increase of purine frequency at 5’ ends was found in animal remains with a more widespread distribution of sample ages (Sawyer et al. 2012). Depurination is inferred from an increase in purines at DNA break points, which has been typically measured as an increase in their absolute frequency (Sawyer et al. 2012). Conversely we think that it is more informative to report relative fold enrichment of purines, which is particularly useful when the levels of enrichment between A and G have to be compared. By measuring absolute purine frequencies it has been shown that A is preferentially found at 5’ end break points in samples younger than 100 years, whereas G is found in samples older than 40,000 years (Sawyer et al. 2012). The enrichment of A in young samples has been attributed to a process independent of depurination probably caused by lysosomal nucleases, which preferentially cut A-T base pairs at the 3’ end of their recognition sites, right after cell death (Sawyer et al. 2012). We think that the A enrichment is an artifact that arises from plotting absolute frequencies instead of relative fold enrichments, which showed no difference neither between A and G in herbarium samples (Figure 2B-C) nor in young animal remains (Sawyer et al. 2012). The apparent enrichment in absolute frequency of G in old animal samples (> 10,000 years) (Sawyer et al. 2012) seems to be a consequence of less A depurination and not more G depurination when fold enrichments are compared. The enrichment in absolute frequency of G has been found even more manifest in a middle Pleistocene horse (Orlando et al. 2011), a phenomenon that holds even when relative fold enrichments are compared. It has been proposed that the increased G enrichment could be the consequence of an electron resonance structure unique to guanosine, which decreases the activation energy to break the N-glycosyl bond making G more labile to depurination (Overballe-Petersen et al. 2012). It is possible that due to the young age of our samples we do not observe the G enrichment at all.

DNA decay and degradation can be understood as a two-step process, with a first rapid phase where the damage is caused mainly by nucleases and digestion by microorganisms, and a second phase where the damage is driven by hydrolytic and oxidative reactions that occur at a much lower rate than the first phase (Molak and Ho 2011). In herbarium samples it is however difficult to assess how different methods of sample preparation, e.g. desiccation, influence the rate and outcome of the first phase. Independent of that, it is possible that the first phase homogenize the differences in degradation between samples of different ages, since historic herbarium and animal samples of very different ages are all highly fragmented. The correlation between fragmentation and time might be the result of a process occurring in the second phase that can be only detected in samples that have experience very similar environments, as it is the case of herbarium samples. The lack of correlation we found between collection year and enrichment in A or G indicates that depurination contributes equally through time to DNA breakage and is to a great extent the underlying process causing the correlation between fragment length and age.

Modern herbarium samples did not show any age-related fragmentation or enrichment of any particular nucleotide at DNA breaking points. As a matter of fact the distribution of fragment lengths was centered on the intended fragment length during sonication (Supplementary Fig. 1). DNA retrieved from our modern herbarium samples was in fact indistinguishable from DNA retrieved from modern tissue. It has been previously suggested based on PCR-based methods that most of DNA fragmentation in herbarium samples occurs during sample desiccation (drying at 60 °C for 18 hours) before they are fixed on herbarium sheets, and only a small portion of damage could be attributed to long-term storage (Staats et al. 2011). We did not find the sample preparation effect in our herbarium samples, however it is worth mentioning that on the contrary to previous studies (Staats et al. 2011) we did not use heating to dry our herbarium samples, since it is well established that heat increases the rate of depurination and subsequently ß elimination leading to DNA strand breaks (Lindahl and Nyberg 1972). Due to the high level of resolution achieved by library-based methods coupled with HTS we could clearly show how fragment length decreases with time, a process which is only manifest in phase two.

### DNA decay rate

DNA fragmentation generates a negative exponential correlation between DNA fragment lengths and the number of amplifiable molecules (Deagle et al. 2006; Schwarz et al. 2009; Adler et al. 2011). This relation allows the calculation of the rate at which bonds are broken in the DNA backbone (lambda) (Deagle et al. 2006). Given a number of dated samples where lambda can be estimated, it is possible to calculate the damage rate of DNA in per site per year units (k). Following this logic it has been recently shown that the temporal decay of DNA in bone follows first-order kinetics and thus can be accurately described by an exponential decay (Allentoft et al. 2012). We have calculated lambda for every herbarium sample based on the distribution of DNA fragment lengths, since the number of DNA fragments decrease exponentially with fragment length (Allentoft et al. 2012) (Fig. 3B). Subsequently we have calculated the DNA decay rate in plant dried tissue using the collection year of the herbarium samples as it has been done with dated animal fossils (Allentoft et al. 2012) (Fig. 3C).

We found that the DNA decay rate in herbarium samples is about ten times faster than the rate in bones (Allentoft et al. 2012) (Figure 4A). It is possible that the big differences in decay rate between herbarium samples and animal remains could be explained by the characteristic nature of each tissue. In bone DNA is absorbed to hydroxyapatite, which decreases the rate of depurination compared to free DNA (Lindahl 1993). Additionally hydroxyapatite binds nucleases (Brundin et al. 2013), which further prevents DNA degradation, especially in the first rapid phase of DNA degradation. DNA in plants’ desiccated tissue might be less protected and more exposed to enzymatic, hydrolytic and oxidative damages. Furthermore, the vast majority of herbarium samples are not mounted on acid-free paper. Acidic paper was regularly used, which could have contributed to DNA degradation, since acid pH increase the rate of depurination in vitro (Lindahl and Nyberg 1972). We calculate independently nuclear and chloroplast DNA decay rates and found that the chloroplast DNA decay rate is 0.75 times the nuclear rate (Fig. 5A). It has been reported that in fossilized bone the mitochondrial DNA decay rate is 2-2.5 times slower than the nuclear one (Allentoft et al. 2012), in agreement with a study that reported a better preservation of mitochondrial relative to nuclear DNA in permafrost mammoth remains (Schwarz et al. 2009). The slower decay rate in organelle DNA might be a consequence of its circular structure, which makes DNA less accessible to endonucleases (Allentoft et al. 2012). An early report of equal rates of degradation between nuclear and chloroplast DNA in herbarium samples has been based in a smaller dataset only interrogated by PCR-based methods, and could be a consequence of lacking experimental resolution (Staats et al. 2011).

We found that the half-life of a 100 bp molecule in a herbarium sample was 40 years (Figure 4B), with a per nucleotide decay rate *k* = 1.66 * 10^−4^ per year. Nevertheless, this estimated half-life is an underestimation, since the extrapolation does not take into account the rapid DNA degradation occurring during the first phase, which is not associated necessarily with time, but could be related to other factors such as the speed of drying of plant tissue (Savolainen et al. 1995). Our half-life calculations predict extremely short fragments in early collected herbarium samples, which make DNA retrieval and sequencing extremely difficult. Fortunately recent advances in DNA extraction (Dabney et al. 2013a) and library preparation (Gansauge and Meyer 2013) developed for middle Pleistocene bones are able to recover very short (< 35 bp) DNA fragments, and could in principle be adapted to extract and sequence DNA from older herbarium specimens.

### DNA misincorporation

In DNA hydrolytic attacks result in modified bases through deamination. These modified bases are misread by polymerases and lead to nucleotide misincorporations. Deamination of C to U leads to the incorporation of A, which will result in C to T or its mirror image G to A substitutions, depending on the sequenced strand. C to T and G to A are the predominant type of substitutions found in aDNA (Hofreiter et al. 2001) and occur principally at the molecule ends as a result of C deamination at single-stranded DNA overhangs (Briggs et al. 2007). We observed also an increase in the percentage of C to T substitutions at the end of the molecule (Fig. 5A) and found a strong correlation between deamination and age (Fig. 5B), as it has been found using animal remains (Sawyer et al. 2012). Although chloroplast reads were less deaminated, the correlation between deamination and age held also for them (Fig. 6B). Notably modern herbarium samples did not show excess of any misincorporation and resembled DNA extracted from fresh tissue.

Since the signal of C deamination has been found recurrently in aDNA studies and there is a strong positive relationship between deamination and sample age (Sawyer et al. 2012), the presence of deamination patterns in aDNA HTS studies has been proposed as authenticity criterion (Krause et al. 2010). It is remarkable that C to T substitutions from both animal remains (Sawyer et al. 2012) and our data correlates strongly with time, although at a different rate in the two tissues, which implies that deamination is strongly related to the phase two of slow DNA degradation. An excess of C to T substitution at the end of the molecule has been also found in plant (Yoshida et al. 2013) and human pathogens (Bos et al. 2011; Schuenemann et al. 2013; Bos et al. 2014) DNA. We found here that the deamination in plant-pathogen-derived reads is intermediate between nuclear- and chloroplast-derived reads (Supplementary Fig. 4). However we think that the signal is sufficient to be used as authenticity criterion. In the future –given an appropriate depth of coverage- it might be possible to also use deamination patterns to authenticate metagenomic aDNA derived from plant or animal tissue, or from environmental DNA profiling.

### Practical implications

It has been suggested that aDNA is a biochemical concept (Shapiro and Hofreiter 2014), which entails that a definition of aDNA does not delineate a time boundary between ancient and modern DNA. We found that DNA retrieved from historic herbarium samples is highly fragmented and contains biochemical changes that lead to misincorporation of nucleotides during sequencing. Therefore, we refer to the DNA extracted from centuries old plant samples as aDNA despite their evolutionarily young age. Modern herbarium samples instead resembled DNA extracted from fresh tissue, which shows that drying by pressing is an ideal method to collect plant samples in long field trips. This also implies that the magnitude of damage that happens in the first phase is highly dependent on the method use to prepare the herbarium specimen.

DNA misincorporations can be confused with natural variation, which will compromise variant calling and increase terminal branches in a phylogenetic context. Both effects are especially prominent in highly deaminated (old) samples that are sequenced at low coverage. Fortunately it is now possible to almost eliminate this source of error by either removing uracils from DNA molecules during library preparation (Briggs et al. 2010), or by statistically distinguishing true variants from aDNA associated misincorporations post-sequencing, in reads derived from single-stranded library preparation methods (Gansauge and Meyer 2013). On the other hand, and due to the strong correlation between accumulation of deaminated residues and time in aDNA, DNA misincorporation remains as one of the most useful criteria to authenticate aDNA reads (Prufer and Meyer 2015).

Unfortunately DNA in dried tissue becomes irremediably shorter through time. Based on our DNA decay rate calculations we predict very short fragments from early 17^th^ century herbarium samples, which will require the use of DNA and library extraction preparation methods capable to recover short length molecules (Dabney et al. 2013a; Gansauge and Meyer 2013). The high DNA fragmentation of historic herbarium samples poses a challenge to genome reduce-representation methods such as RAD (restriction site associated markers)-sequencing (Miller et al. 2007; Baird et al. 2008). Short DNA fragments will have in theory less restriction sites to offer to restriction enzymes. Additionally the use of restriction enzymes will reduce further the length of already short DNA fragments, which will cause that a big fraction of the reads cannot be mapped to a reference genome due to its very short size. The use of RAD-sequencing to DNA retrieved from museum specimens has shown low DNA yields and low percentage of reads that could be mapped to the reference genome (Tin et al. 2014). Another limitation of short fragment lengths is the difficulty it adds in assembling ancient genomes *de novo*. Thanks to improvements in library preparation and HTS accuracy, it is however possible to sequence and perform mapping-guided assemblies of complete genomes from historic specimens with quality that matches genomes derived from modern specimens and therefore exploit the millions of plant remains stored in herbaria worldwide.

## Material and methods

### Previously published datasets

Sequences derived from *Solanum tuberosum* and *Solanum lycopersicum* infected by *Phytophthora infestans* are deposited in the European Nucleotide Archive, with accession number PRJEB1877.

### Herbarium samples

Historic herbarium samples were either directly sampled by us in different herbaria both in North America and Europe, or sampled there by collection curators and sent to us by post (Supplementary Table S1). The amount of tissue used for destructive sampling ranged from 2 to 8 mm^2^.

Modern herbarium samples derived from a recent collection of *A. thaliana* wild populations in North America by the Max Planck Institute for Developmental Biology. After collection plant tissue was dried by pressing between acid-free papers using a wooden press for four 6 weeks and subsequently mounted in herbarium sheets.

### DNA extraction, library preparation and sequencing

DNA extraction from historic herbarium samples: DNA extractions were carried out in clean room facilities in all cases. The majority of the samples were extracted following the PTB extraction protocol (Kistler 2012) as previously described (Yoshida et al. 2013). Samples from the Cornell Bailey Hortorium were extracted using the CTAB extraction protocol (Kistler 2012) (Supplementary Table S2).

DNA extraction of modern herbarium samples: DNA extractions were carried out following the PTB extraction protocol (Kistler 2012).

Library preparation historic samples: Illumina double indexed sequencing libraries (Kircher et al. 2012; Meyer et al. 2012) were prepared from each sample as previously described (Yoshida et al. 2013). The excess of C-to-T substitutions associated with DNA damage and caused by deamination of cytosines (Hofreiter et al. 2001) was not repaired in order to quantify the amount of damage present in samples of different ages.

Library preparation modern herbarium samples: indexed libraries were prepared using the Illumina TruSeq Nano DNA sample preparation kit following manufacturer instructions.

Sequencing: Libraries were paired-end sequenced on the Illumina HiSeq 2000, HiSeq 2500 or MiSeq instruments (Supplementary Table S2).

### Read processing and mapping

Historic herbarium samples: reads were assigned to each sample based on their indices. Adapters were trimmed using the program Skewer (version 0.1.120) with default settings with the natively implemented Illumina TruSeq adapter sequences (Jiang et al. 2014). Forward and reverse reads were merged using the program Flash (version 1.2.11) with default settings, except for an elevated maximum overlap (100-150 bp depending on read length) to allow a more accurate scoring of highly overlapping read pairs (Magoc and Salzberg 2011). Merged reads were mapped as single-end reads to their respective reference genomes: *Arabidopsis thaliana* (Arabidopsis Genome 2000; Swarbreck et al. 2008), *Solanum tuberosum* (Potato Genome Sequencing Consortium, 2011), *Solanum lycopersicum* (Consortium 2012), *Phytophthora infestans* (Haas et al. 2009). The mapping was performed using BWA-MEM (version 0.7.10) with default settings (Li 2013). PCR-duplicates were identified after mapping based on start and end coordinates and for every cluster of duplicate reads a consensus sequence was generated (Kircher 2012).

Modern herbarium samples: reads were processed very similar to the reads that belong to historic samples. The vast majority of reads could not be merged, which indicates that the DNA was not as fragmented as in older herbarium samples. Therefore, we mapped the paired-end reads using BWA-MEM (version 0.7.10) with default parameters (Li 2013) and inferred fragment size based on paired-end mapping.

### Analysis of DNA damage patterns

Fragment length: To characterize the patterns of DNA fragmentation we analyzed the fragment length distributions of merged reads. We fitted a log-Normal distribution to the empirical fragment length distributions using the fitdistr function from the package MASS using R. Since in a log-Normal distribution the logarithm of a variable is normally distributed, we used the mean of this distribution (log-mean) to summarize the fragment length distribution. The regression on the relationship among log-mean of fragment lengths and collection year was carried out using the lm function in R. For visualization (Fig. 1B) we used the fragment length median on a log-scaled y-axis, since the median is more intuitive to understand than the log-mean value. The relationship between log-mean and median follows the formula: *median = e^log-mean^*.

DNA break points: To analyze the nucleotide genomic context around DNA break points we used the software mapDamage 2.0 (version 2.0.2-12) (Jonsson et al. 2013). MapDamage calculates the genomic base frequencies around mapped reads and within reads, which allows the inference of the bases most likely to be present before DNA break points. We calculated the relative enrichment of either adenine or guanine at 5’ end (position -1 compared with position - 5). The frequencies of both adenine and guanine were extracted from the output file dnacomp.txt produced by mapDamage. The regression on the relationship among purine relative enrichment (either adenine or guanine) and collection year was carried out using the lm function in R. The whole procedure was carried out for plant nuclear and chloroplast reads independently.

Nucleotide misincorporation: all types of nucleotide misincorporations relative to the reference genome were calculated per library using mapDamage 2.0 (version 2.0.2-12) (Jonsson et al. 2013). The percentage of C to T substitutions at first base was extracted from the output file 5pCtoT_freq.txt produced by mapDamage. The regression on the relationship among the percentage of C to T substitutions at first base (5’ end) and collection year was carried out using the lm function in R. For the regression we used the percentage of deamination at first base. The whole procedure was carried out for plant nuclear and chloroplast reads independently, and also for pathogen nuclear reads in the case of samples infected with *P. infestans*.

### Calculation of DNA decay rate

To calculate the decay rate of DNA retrieved from plant desiccated tissue we used a previously described methodology (Allentoft et al. 2012) and adapted it to multiple samples. The random fragmentation of DNA molecules that occurs *post mortem* follows a model of exponential decay, i.e. the amount amplifiable template decreases exponentially with increasing length (Deagle et al. 2006). We used the distributions of fragment length (*L*) of mapped reads to calculate the DNA decay rate, which is determined by the proportion of damage sites (*λ*). Thus, the process can be described using an exponential distribution:

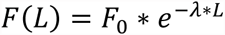

Where *L* is the fragment length, *F(L)* the frequency of fragment with length *L* and *F*_*O*_ the frequency intersect at length 0.

After logarithmic transformation there is a linear relationship between the logarithms of the fragment frequency and fragment length with a slope *– λ*:

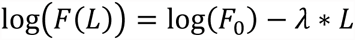

In this relationship *λ* describes the fraction of bond survival per base in a single sample/library (Deagle et al. 2006; Allentoft et al. 2012). As previously described (Allentoft et al. 2012), the DNA decay rate per base per year *k* can then be calculated as:

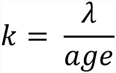

We calculated the decay rate across all analyzed samples taking advantage of the negative correlation between fragment length and age of the sample. We plotted the damage fraction per site *(λ)* as a function of sample age. The slope of the linear regression on the relationship among *λ* and samples age yields *k*, the decay rate, according to the linear relationship:

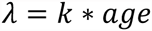

The whole procedure was carried out for plant nuclear and chloroplast reads independently.

### Analysis of covariance

To test if the regressions between chloroplast- and nuclear-derived reads were significantly different we performed an analysis of covariance. We used the “aov” function in R to test models where the sample age was the covariate and the DNA origin (chloroplast- or nuclear-derived) was the factor. In the first step, a model of type “y ∼ covariate * factor” was used to include a possible interaction between covariate and factor, which would mean that there is a difference in the slope of the regression depending on the factor. If no significant interaction was detected, the “anova” command in R was used to test this model against a model of type “y ∼ covariate + factor”. This last model does not include the interaction, therefore we can test whether the removal of the interaction has an effect on the fit of the model. If not, the second model was accepted with the conclusion, that the regressions do not differ in slope, but possibly in their intersects (if there is a significant effect of the factor on the dependent variable y).

To test whether the linear regressions of read lengths and collection year between samples extracted with CTAB and PTB methods were different, we used the same approach as for chloroplast- and nuclear-derived reads. In this comparison we used extraction method as the factor in the linear model.

### Data access

New DNA sequences are deposited in the European Nucleotide Archive, with accession number PRJEB9878.

## Acknowledgements

We thank Moisés Espósito-Alonso, Rafal Gutaker and Rebecca Schwab for helpful comments on the manuscript; Matthias Meyer and Petra Korlević for input on data analysis and guidance through specialized literature; Michael Dannemann for statistical advice; Magdalena Grenda for input on herbaria conservation. We are grateful to all researchers and curators for providing the historic specimens: Marco Thines (Goethe Universität, Frankfurt, Germany), Sandra Knapp (Natural History Museum, London, UK), Bryn Dentinger (Royal Botanical Gardens, Kew, UK), Dagmar Triebel (Botanishe Staatssammlung, Munich, Germany), Anna Stalter (Cornell Bailey Hortorium, Ithaca, USA), John Freudenstein (The Ohio State University, Columbus, USA), Christine Niezgoda (The Field Museum, Chicago, USA), Robert Capers (University of Connecticut, Storrs, USA), Carol Ann McCormick (University of North Carolina, Chapel Hill, USA), Michael S. Dossmann (Arnold Arboretum, Boston, USA), Cathy M. Herring (Agricultural Research Station, North Carolina State University, Clayton, USA), John Peter (New York Botanical Garden, New York, USA). We thank Oliver Haddrath for laboratory space for DNA extraction in the Royal Ontario Museum. This study was funded by the Presidential Innovation Fund of the Max Planck Society.

### Disclosure declaration

The authors declare that no competing interests exist.

### Author contributions

C.L.W. and H.A.B conceived and designed the experiments. V.J.S, E.R, G.S., J.D., B.A.G. performed DNA extractions; V.J.S., E.R, J.D. performed libraries; J.R.S. and J.K. coordinated laboratory experiments; C.L.W. and H.A.B. analyzed the data; C.L.W and H.A.B wrote the paper with help from all authors.

**Figure S1.**
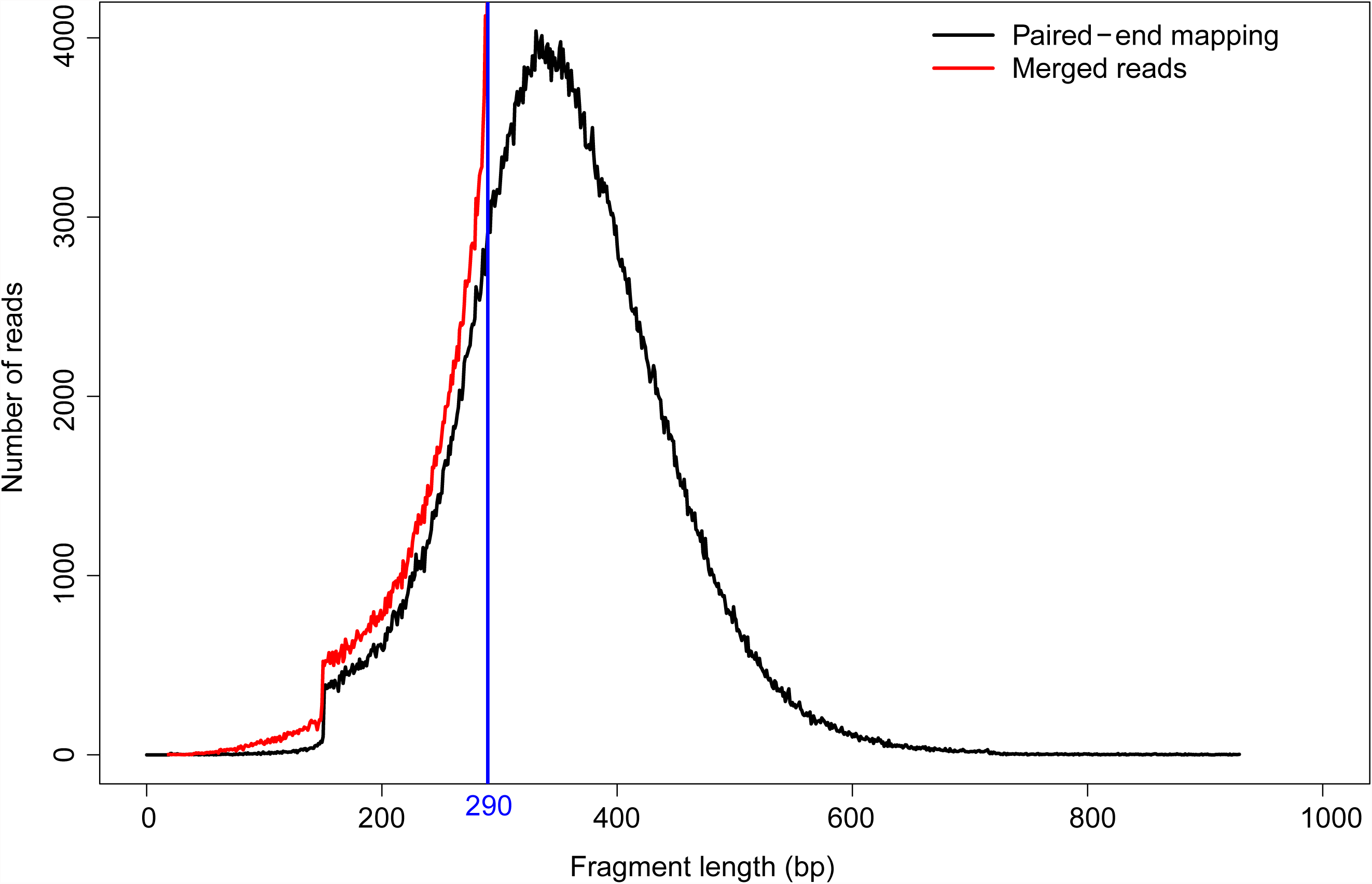
DNA fragmentation in recently prepared herbarium samples. The black line shows the fragment length distribution inferred after performing paired-end mapping. The red line shows the fragment length distribution for reads that could be merged. The horizontal blue line indicates the maximum length of merged reads from 2×150 bp paired-end reads (290 bp), when a minimum overlap of 10 bp is required between forward and reverse reads.

**Figure S2.**
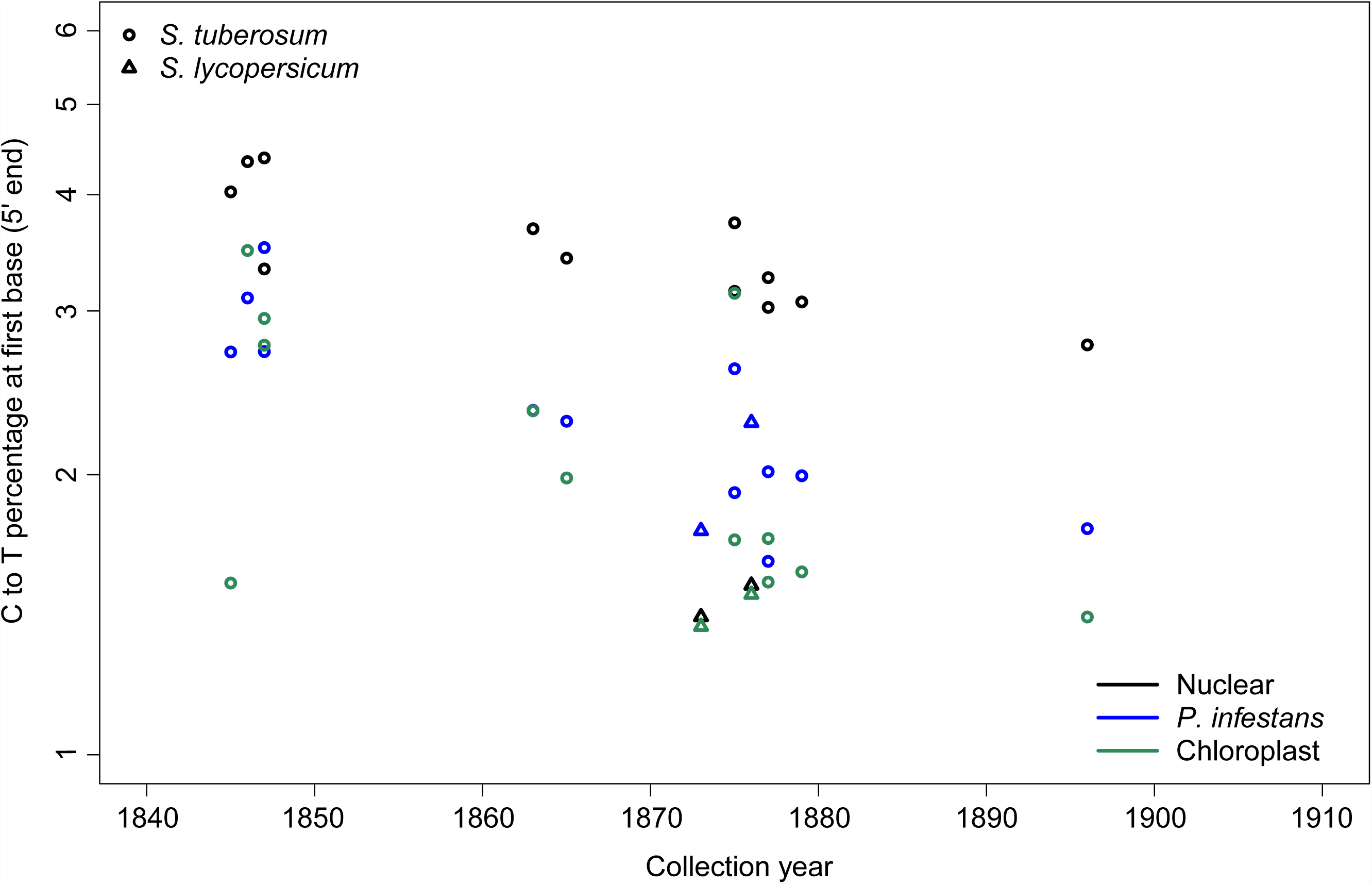
Nucleotide misincorporation in samples with lesions compatible with *Phytophthora infestans*. C to T percentage at first base (5′ end) as a function of collection year for host and pathogen reads. Host reads are split in nuclear and chloroplast reads.

**Figure S3.**
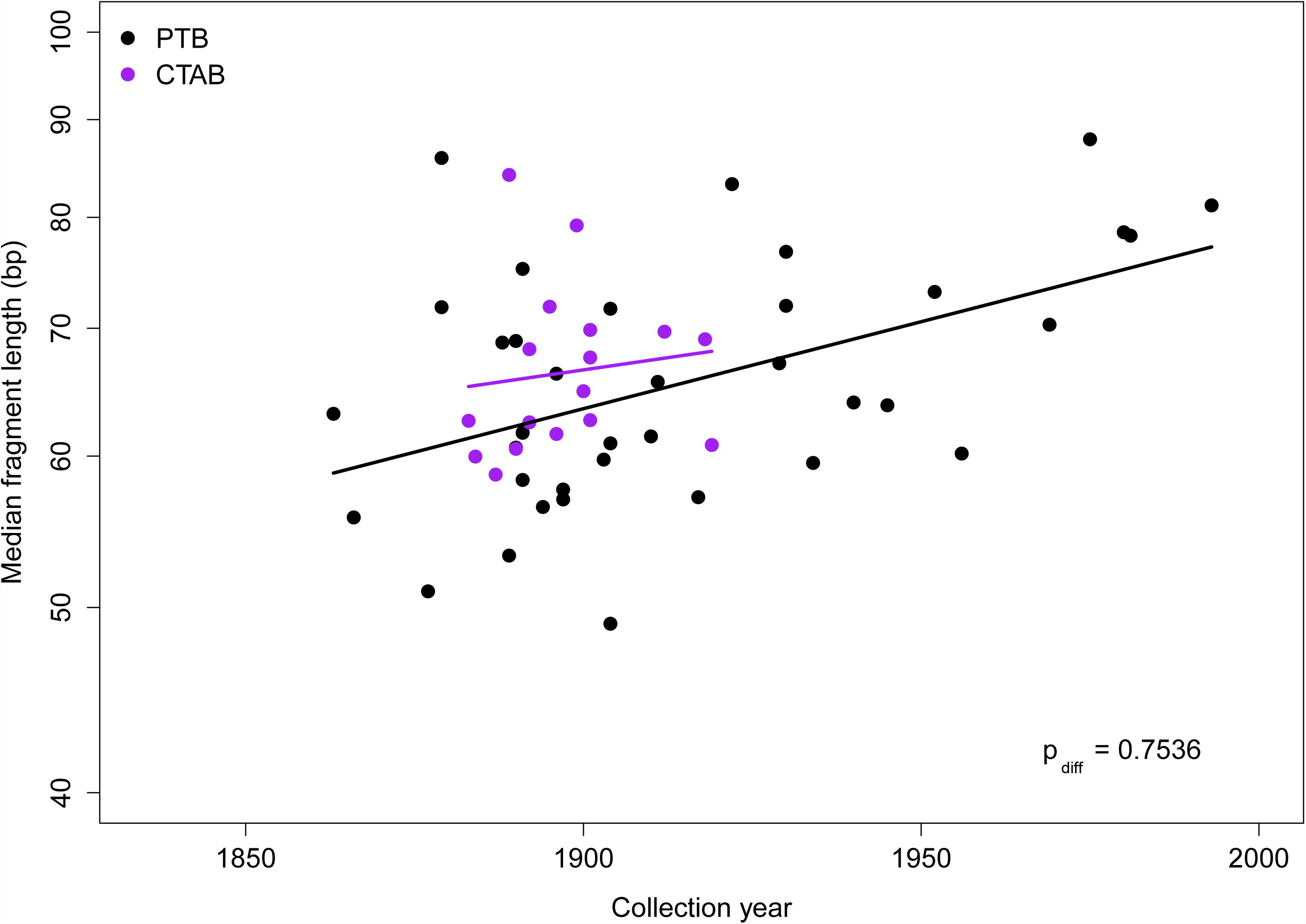
DNA fragmentation of *A. thaliana* samples extracted using CTAB and PTB method. Median length of merged reads from both methods as a function of collection year. The lines indicates the linear regression

**Figure S4.**
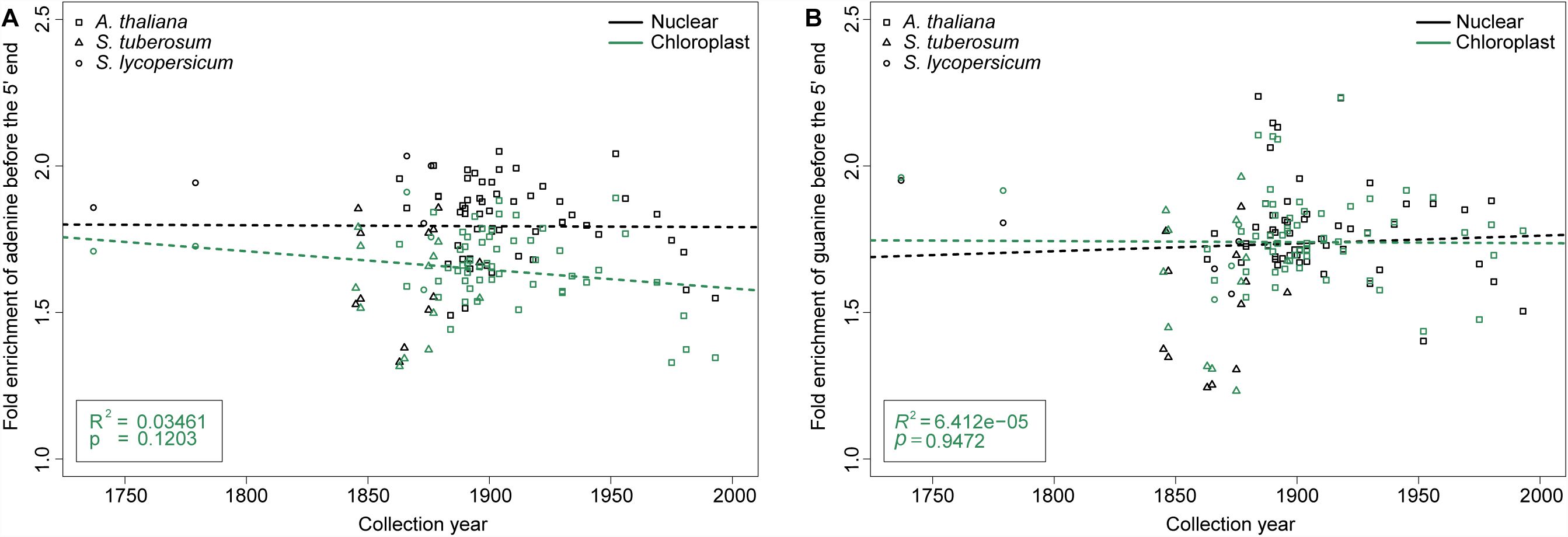
DNA breaking points in nuclear and chloroplast reads. **(A)** Relative enrichment of adenine at 5’ end (position -1 compared with position -5) as a function of collection year **(B)** Relative enrichment of guanine at 5’ end (position -1 compared with position -5) as a function of collection year. In both (A) and (B) data points, regression lines and values for nuclear and chloroplast reads are shown in black and green font, respectively.

---

**Table S1.** Provenance of herbaria samples.

| **ID** | **Country of origin** | **Collection year** | **Species** | **Reference** |
| --- | --- | --- | --- | --- |
| BH0000052681 | USA | 1919 | *A. thaliana* | 1 |
| BH0000061407 | USA | 1890 | *A. thaliana* | 1 |
| BH0000061409 | USA | 1892 | *A. thaliana* | 1 |
| BH0000061416 | USA | 1892 | *A. thaliana* | 1 |
| BH0000061457 | USA | 1900 | *A. thaliana* | 1 |
| BH0000061454 | USA | 1884 | *A. thaliana* | 1 |
| BH0000061397 | USA | 1899 | *A. thaliana* | 1 |
| BH0000061417 | USA | 1912 | *A. thaliana* | 1 |
| BH0000061449 | USA | 1901 | *A. thaliana* | 1 |
| BH0000061420 | USA | 1896 | *A. thaliana* | 1 |
| BH0000061395 | USA | 1883 | *A. thaliana* | 1 |
| BH0000061424 | USA | 1895 | *A. thaliana* | 1 |
| BH0000061393 | USA | 1918 | *A. thaliana* | 1 |
| BH0000061399 | USA | 1901 | *A. thaliana* | 1 |
| BH0000061448 | USA | 1901 | *A. thaliana* | 1 |
| BH0000061418 | USA | 1887 | *A. thaliana* | 1 |
| BH0000061406 | USA | 1889 | *A. thaliana* | 1 |
| OSU46488 | USA | 1917 | *A. thaliana* | 2 |
| OSU13896 | USA | 1930 | *A. thaliana* | 2 |
| OSU13900 | USA | 1934 | *A. thaliana* | 2 |
| OSU54623 | USA | 1956 | *A. thaliana* | 2 |
| OSU79944 | USA | 1911 | *A. thaliana* | 2 |
| OSU183663 | USA | 1969 | *A. thaliana* | 2 |
| OSU361885 | USA | 1930 | *A. thaliana* | 2 |
| OSU150287 | USA | 1980 | *A. thaliana* | 2 |
| OSU163761 | USA | 1981 | *A. thaliana* | 2 |
| OSU364632 | USA | 1993 | *A. thaliana* | 2 |
| UNC54051 | USA | 1945 | *A. thaliana* | 3 |
| UNC25707 | USA | 1940 | *A. thaliana* | 3 |
| UNC63978 | USA | 1910 | *A. thaliana* | 3 |
| CT79391 | USA | 1904 | *A. thaliana* | 4 |
| CT79409 | USA | 1929 | *A. thaliana* | 4 |
| CT79389 | USA | 1975 | *A. thaliana* | 4 |
| 176849CFM | USA | 1904 | *A. thaliana* | 5 |
| 531679CFM | USA | 1922 | *A. thaliana* | 5 |
| 1507461CFM | USA | 1952 | *A. thaliana* | 5 |
| NY102365 | USA | 1903 | *A. thaliana* | 6 |
| NY1365344 | USA | 1890 | *A. thaliana* | 6 |
| NY888124 | USA | 1863 | *A. thaliana* | 6 |
| NY1365363 | USA | 1888 | *A. thaliana* | 6 |
| NY888144 | USA | 1866 | *A. thaliana* | 6 |
| NY1365364 | USA | 1889 | *A. thaliana* | 6 |
| NY888134 | USA | 1877 | *A. thaliana* | 6 |
| NY1365332 | USA | 1890 | *A. thaliana* | 6 |
| NY1365337 | USA | 1891 | *A. thaliana* | 6 |
| NY1365370 | USA | 1897 | *A. thaliana* | 6 |
| NY1365333 | USA | 1894 | *A. thaliana* | 6 |
| NY1365374 | USA | 1896 | *A. thaliana* | 6 |
| NY102364 | USA | 1904 | *A. thaliana* | 6 |
| NY1365375 | USA | 1897 | *A. thaliana* | 6 |
| NY888141 | USA | 1879 | *A. thaliana* | 6 |
| NY1365354 | USA | 1891 | *A. thaliana* | 6 |
| NY888141 | USA | 1879 | *A. thaliana* | 6 |
| NY1365354 | USA | 1891 | *A. thaliana* | 6 |
| KM177509 | England | 1865 | *S. tuberosum* | 7 |
| KM177517 | Wales | 1875 | *S. tuberosum* | 7 |
| KM177514 | Ireland | 1847 | *S. tuberosum* | 7 |
| KM177500 | England | 1845 | *S. tuberosum* | 7 |
| KM177513 | Ireland | 1846 | *S. tuberosum* | 7 |
| KM177548 | England | 1847 | *S. tuberosum* | 7 |
| M-0182898 | Germany | 1863 | *S. tuberosum* | 8 |
| M-0182906 | Germany | 1877 | *S. tuberosum* | 8 |
| M-0182907 | Germany | 1875 | *S. tuberosum* | 8 |
| M-0182896 | Germany | 1877 | *S. tuberosum* | 8 |
| M-0182903 | Canada | 1896 | *S. tuberosum* | 8 |
| M-0182904 | Austria | 1879 | *S. tuberosum* | 8 |
| BM000777791 | UK | 1866 | *S. lycopersicum* | 9 |
| BM000815937 | UK | 1737 | *S. lycopersicum* | 9 |
| BM000849510 | UK | 1779 | *S. lycopersicum* | 9 |
| M-0182897 | USA | 1876 | *S. lycopersicum* | 8 |
| M-0182900 | Germany | 1873 | *S. lycopersicum* | 8 |
| *Modern herbarium* |  |  |  |  |
| INTHRR_108 | USA | 2014 | *A. thaliana* | 10 |
| INTHRR_110 | USA | 2014 | *A. thaliana* | 10 |
| INTHRR_54 | USA | 2014 | *A. thaliana* | 10 |
| MAAA_16 | USA | 2014 | *A. thaliana* | 10 |
| MAAA_41 | USA | 2014 | *A. thaliana* | 10 |
| MAAA_61 | USA | 2014 | *A. thaliana* | 10 |
| MDTCR_32 | USA | 2014 | *A. thaliana* | 10 |
| MDTCR_42 | USA | 2014 | *A. thaliana* | 10 |
| MDTCR_51 | USA | 2014 | *A. thaliana* | 10 |
| NCARS_32 | USA | 2014 | *A. thaliana* | 10 |
| NCARS_51 | USA | 2014 | *A. thaliana* | 10 |
| NCARS_9 | USA | 2014 | *A. thaliana* | 10 |
| NYBG_154 | USA | 2014 | *A. thaliana* | 10 |
| NYBG_157 | USA | 2014 | *A. thaliana* | 10 |
| NYBG_169 | USA | 2014 | *A. thaliana* | 10 |

\* 1. Cornell Bailey Hortorium; 2. Ohio State University Herbarium; 3. University of North Carolina Herbarium; 4. University of Connecticut Herbarium; 5. Chicago Field Museum; 6. New York Botanical Garden; 7. Kew Royal Botanical Garden; 8. Botanische Staatssammlung München; 9. Natural History Museum, London; 10. Max Planck Institute for Developmental Biology collection.

**Table S2.** Sequencing strategy and summary statistics.

| **ID** | **Read type** | **Number raw read pairs** | **Number merged reads** | **Percentage of reads mapped to reference** |
| --- | --- | --- | --- | --- |
| BH0000052681 | 2x101 | 4,070,122 | 3,995,427 | 53.5 |
| BH0000061407 | 2x101 | 4,083,714 | 4,035,185 | 28.4 |
| BH0000061409 | 2x101 | 4,316,387 | 4,058,032 | 66.5 |
| BH0000061416 | 2x101 | 3,927,269 | 3,857,422 | 71.3 |
| BH0000061457 | 2x101 | 478,438 | 467,456 | 74.6 |
| BH0000061454 | 2x101 | 497,373 | 491,061 | 66.6 |
| BH0000061397 | 2x101 | 589,715 | 542,051 | 89.5 |
| BH0000061417 | 2x101 | 3,310,517 | 3,243,814 | 76.9 |
| BH0000061449 | 2x101 | 4,873,199 | 4,812,870 | 92.4 |
| BH0000061420 | 2x101 | 4,137,492 | 4,051,781 | 81.0 |
| BH0000061395 | 2x101 | 608,442 | 593,440 | 87.6 |
| BH0000061424 | 2x101 | 560,391 | 546,601 | 83.4 |
| BH0000061393 | 2x101 | 583,208 | 567,480 | 81.7 |
| BH0000061399 | 2x101 | 5,030,578 | 4,882,226 | 88.5 |
| BH0000061448 | 2x101 | 3,250,580 | 3,179,443 | 60.9 |
| BH0000061418 | 2x101 | 4,110,260 | 4,063,919 | 72.3 |
| BH0000061406 | 2x101 | 5,525,374 | 5,236,914 | 85.2 |
| OSU46488 | 2x150 | 775,962 | 732,837 | 69.3 |
| OSU13896 | 2x150 | 1,222,514 | 1,157,217 | 73.5 |
| OSU13900 | 2x150 | 885,259 | 822,341 | 42.0 |
| OSU54623 | 2x150 | 814,706 | 768,157 | 69.1 |
| OSU79944 | 2x150 | 935,769 | 883,902 | 76.5 |
| OSU183663 | 2x150 | 941,643 | 894,825 | 68.6 |
| OSU361885 | 2x150 | 815,261 | 770,652 | 74.9 |
| OSU150287 | 2x150 | 1,100,356 | 1,025,252 | 68.8 |
| OSU163761 | 2x150 | 1,026,456 | 982,279 | 85.0 |
| OSU364632 | 2x150 | 992,421 | 930,033 | 68.7 |
| UNC54051 | 2x150 | 998,331 | 923,388 | 86.7 |
| UNC25707 | 2x150 | 741,049 | 645,792 | 71.6 |
| UNC63978 | 2x150 | 987,305 | 936,195 | 74.4 |
| CT79391 | 2x150 | 928,321 | 886,829 | 70.7 |
| CT79409 | 2x150 | 854,432 | 802,723 | 81.8 |
| CT79389 | 2x150 | 781,485 | 745,982 | 30.4 |
| 176849CFM | 2x150 | 918,792 | 874,775 | 61.5 |
| 531679CFM | 2x150 | 870,119 | 809,957 | 83.2 |
| 1507461CFM | 2x150 | 924,012 | 877,358 | 78.4 |
| NY102365 | 2x150 | 1,313,215 | 1,269,942 | 67.9 |
| NY1365344 | 2x150 | 1,496,570 | 1,432,778 | 80.6 |
| NY888124 | 2x150 | 2,083,306 | 2,009,800 | 70.0 |
| NY1365363 | 2x150 | 1,408,827 | 1,354,608 | 74.9 |
| NY888144 | 2x150 | 1,664,389 | 1,592,414 | 64.6 |
| NY1365364 | 2x150 | 1,660,257 | 1,597,623 | 60.3 |
| NY888134 | 2x150 | 1,333,231 | 1,279,207 | 75.9 |
| NY1365332 | 2x150 | 2,093,699 | 2,013,728 | 80.5 |
| NY1365337 | 2x150 | 1,773,246 | 1,709,015 | 81.3 |
| NY1365370 | 2x150 | 1,322,896 | 1,282,512 | 80.1 |
| NY1365333 | 2x150 | 1,937,923 | 1,864,708 | 79.4 |
| NY1365374 | 2x150 | 1,785,933 | 1,727,387 | 75.9 |
| NY102364 | 2x150 | 1,621,005 | 1,561,419 | 59.2 |
| NY1365375 | 2x150 | 1,713,640 | 1,638,994 | 36.3 |
| NY888141 | 2x101 | 12,574,596 | 12,286,269 | 83.9 |
| NY1365354 | 2x101 | 10,832,427 | 10,602,196 | 66.5 |
| NY888141 | 2x101 | 9,605,582 | 9,485,526 | 86.1 |
| NY1365354 | 2x101 | 11,982,636 | 11,796,667 | 78.9 |
| KM177509 | 2x151 | 666,206 | 662,804 | 70.1 |
| KM177517 | 2x151 | 1,017,778 | 1,013,025 | 62.6 |
| KM177514 | 2x151 | 803,024 | 798,164 | 53.5 |
| KM177500 | 2x151 | 808,056 | 802,648 | 30.8 |
| KM177513 | 2x151 | 704,372 | 702,024 | 60.0 |
| KM177548 | 2x151 | 611,321 | 609,290 | 53.9 |
| M-0182898 | 2x101 | 2,206,547 | 1,843,207 | 65.0 |
| M-0182906 | 2x101 | 2,441,550 | 2,313,611 | 75.6 |
| M-0182907 | 2x101 | 2,224,182 | 2,100,559 | 75.4 |
| M-0182896 | 2x101 | 2,735,620 | 2,506,508 | 55.1 |
| M-0182903 | 2x101 | 3,668,118 | 3,461,916 | 65.3 |
| M-0182904 | 2x101 | 2,989,144 | 2,855,446 | 83.7 |
| BM000777791 | 2x101 | 533,872 | 518,134 | 91.9 |
| BM000815937 | 2x101 | 554,635 | 551,331 | 66.1 |
| BM000849510 | 2x101 | 519,254 | 515,461 | 68.9 |
| M-0182897 | 2x101 | 3,775,244 | 3,624,340 | 79.7 |
| M-0182900 | 2x101 | 1,860,985 | 1,582,951 | 85.7 |
| Modern Herbarium |  |  |  |  |
| INTHRR_108 | 2x150 | 1,013,210 | 173,455 | 85.5 |
| INTHRR_110 | 2x150 | 614,641 | 107,820 | 76.6 |
| INTHRR_54 | 2x150 | 1,035,098 | 183,394 | 80.8 |
| MAAA_16 | 2x150 | 816,998 | 151,363 | 90.3 |
| MAAA_41 | 2x150 | 994,861 | 196,576 | 68.0 |
| MAAA_61 | 2x150 | 1,029,669 | 192,361 | 90.0 |
| MDTCR_32 | 2x150 | 805,969 | 127,286 | 87.0 |
| MDTCR_42 | 2x150 | 977,367 | 206,661 | 91.5 |
| MDTCR_51 | 2x150 | 175,376 | 35,842 | 90.7 |
| NCARS_32 | 2x150 | 748,794 | 141,618 | 86.8 |
| NCARS_51 | 2x150 | 949,138 | 216,536 | 88.8 |
| NCARS_9 | 2x150 | 787,902 | 139,538 | 85.0 |
| NYBG_154 | 2x150 | 1,867,301 | 482,893 | 68.1 |
| NYBG_157 | 2x150 | 2,001,726 | 801,146 | 62.0 |
| NYBG_169 | 2x150 | 1,691,752 | 504,633 | 75.8 |

